# Engineering an RNA-based tissue-specific platform for genetic editing through use of a miRNA-enabled Cas12a

**DOI:** 10.1101/2020.03.04.977363

**Authors:** Rasmus Møller, Kohei Oishi, Benjamin R. tenOever

## Abstract

The capacity to edit genomes using CRISPR-Cas systems holds immense potential for countless genetic-based diseases. However, one significant impediment preventing broad therapeutic utilization is *in vivo* delivery. While genetic editing at a single cell level *in vitro* can be achieved with high efficiency, the capacity to utilize these same biologic tools in a desired tissue *in vivo* remains challenging. Non-integrating RNA virus-based vectors constitute efficient platforms for transgene expression and surpass several barriers to *in vivo* delivery. However, the broad tissue tropism of viral vectors raises the concern for off-target effects. Moreover, prolonged expression of the Cas proteins, regardless of delivery method, can accumulate aberrant RNAs leading to unwanted immunological responses. In an effort to circumvent these shortcomings, here we describe a versatile RNA virus-based technology that can achieve cell-specific activity and self-inactivation by combining host microRNA (miRNA) biology with the CRISPR-Cas12a RNA-guided nuclease. Exploiting the RNase activity of Cas12a, we generated a vector that self-inactivates upon delivery of Cas12a and an accompanying CRISPR RNA (crRNA). Furthermore, we show that maturation of the crRNA can be made dependent on cell-specific miRNAs, which confers cell-specificity. We demonstrate that this genetic editing circuit delivers diminished yet sufficient levels of Cas12a to achieve effective genome editing whilst inducing a minimal immunological response. It can also function in a cell-specific manner thereby facilitating *in vivo* editing and mitigating the risk of unwanted, off-target effects.

## INTRODUCTION

The CRISPR-Cas systems of archaea and many bacteria are sequence-specific adaptive defense systems that have evolved to cleave foreign nucleic acid^1^. This defense system is dependent on acquisition and integration of foreign DNA protospacers in a process generally referred to as adaptation^2^. Once integrated, expression of the so-called protospacers generate a precursor CRISPR RNA (pre-crRNA) which is further processed and matured to produce crRNA – comprised of a 5’ direct repeat and the spacer. Finally, crRNA is bound by a Cas nuclease to elicit interference on incoming DNA as defined by complementarity of its guide RNA. Moreover, as the protospacer DNA is inherited, adaptation of a single prokaryotic cell can result in Lamarckian evolution for its offspring^3^.

While most of the Cas-nucleases only possess RNA-guided DNase activity, Cas12a also has RNase function^4^. The RNase function is responsible for processing the pre-crRNA by cleaving direct repeat sequences that flank the protospacer^4^. The crRNA that is generated as a result of these processing events is sufficient for instilling specificity onto the DNase activity of Cas12a. Similar to Cas9, Cas12a has also been repurposed as a eukaryotic gene editor^5–8^. However, as Cas12a biology is still in its infancy, its optimization lags behind that of Cas9. Despite this, the ability of Cas12a to process its own crRNA enables one to use it to generate the crRNA from diverse types of RNA so long as it is flanked by direct repeats^9^. This activity not only allows one to generate multiple crRNAs for any number of targets, but it has also enabled the generation of mRNAs that both code for Cas12a and the desired guides on a single transcript^10^. This is in contrast to the most commonly applied Cas9 editing tool which demands a separate DNA dependent RNA polymerase for production of Cas9 and the single guide RNA^6,11^.

Despite the immense potential of both the Cas9 and Cas12a systems, one significant impediment remains delivery of these large proteins alongside the desired crRNA(s). This challenge is formidable, especially when one wishes to efficiently edit a large number of cells to repair a genetic defect *in vivo*. This problem is further confounded by the fact that maintaining Cas expression for longer periods of time can result in the generation of off-target effects, aberrant RNAs, chromosomal translocations, and/or removal of the Cas-expressing cells^12,13^. Given these challenges, the most optimal genetic editors would be delivered with the efficiency of a virus in a manner that was free of any possibility of genomic integration and would function only in a desired cell type for the time required to achieve editing. To this end, we designed a self-inactivating RNA that expresses Cas12a and processes its own crRNA in a cell-specific manner that is amenable to RNA virus-based delivery.

Having some parallels with the CRISPR-Cas platform, host miRNAs are rooted in defense systems albeit optimized for the targeting of RNA, rather than DNA^14^. Generally described as RNA interference (RNAi), this defense system acquires double stranded RNA (dsRNA) fragments from incoming virus and processes them into small interfering RNAs (siRNAs) similar in length than that of crRNAs. Furthermore, like crRNAs, siRNAs provide specificity to an otherwise non-specific nuclease. In the case of RNAi, siRNAs are bound by a family of Argonaute (AGO) proteins with the capacity to cleave target mRNA in a sequence-specific manner^15^. The emergence of this system was so evolutionarily successful, that the genes involved in this biology duplicated and generated a similar small RNA based system commonly referred to as microRNAs (miRNAs)^16^. Unlike the small RNA-based antiviral systems, miRNAs derive from the host genome but are ultimately processed and utilized in a similar function, even sharing some overlap with the processing machinery of the Type II CRISPR-Cas system^17^. The general repurposing of this biology is believed to have happened independently in plants and animals but, in both examples, the biogenesis of miRNA-mediated regulation enabled another level of transcriptional control which coincided with multicellularity^16^.

One significant difference between antiviral RNAi and host miRNA biology is the complementarity between the small RNA and its target(s). In contrast to antiviral RNAi, miRNAs do not generally bind with perfect complementarity to their targets and therefore do not induce cleavage, owing to the fact that they are derived from host genes^16^. As a result, miRNAs are often considered ‘fine-tuners’ of host biology and are thought to act over the course of days or weeks^18^. In contrast, virus-derived siRNAs can engage their target with complete complementarity, as they derive from the pathogen itself, and thus enable both cleavage and the recycling of the small RNA – achieving silencing within hours^16^. However, as this difference in biology is almost completely defined by complementarity, if one introduces a perfect binding site for a given miRNA into a transcript, it will undergo potent silencing more akin to the normal activity of antiviral RNAi^19^. This exploitation of miRNA biology therefore enables one to generate viruses or virus-based vectors which can be ubiquitously targeted or function in a species-, tissue-, cell-, or even signaling-specifc manner based on endogenous miRNA expression^20–26^.

In an effort to generate an RNA-based DNA editor that functions in a cell-specific manner that would be amenable for *in vivo* use, we combined CRISPR-Cas and miRNA biology. In brief, we utilize the fact that Cas12a processes its own pre-crRNA to make a vector that delivers both Cas12a and crRNA and in doing so, inactivates itself. To this end, we encode crRNAs in the 3’-UTR of Cas12a and show that it leads to self-cleavage of its own transcript. Moreover, we demonstrate that delivery of this self-inactivating construct is sufficient to achieve efficient gene editing. And lastly, we show that processing of the pre-crRNA can be made to be dependent on miRNA-expression thereby conferring cell-type specificity on our editing platform.

## RESULTS

In an effort to build a single mRNA transcript that could both self-inactivate and function in a cell-specific manner, we designed a construct encoding an enhanced green fluorescent protein (GFP) and an HA epitope-tagged Cas12a separated by a P2A peptide site^27^ with its targeting instructions in the 3’ UTR (Fig. 1a). To achieve self-inactivation, we incorporated a crRNA in the 3’ untranslated region (UTR) of the GFP-Cas12a mRNA that consisted of a direct repeat and a spacer (DR). Accurate processing of this crRNA relies on a second motif found 3’ of the spacer comprising a perfect miRNA target (miR-T) site. In the presence of the cognate miRNA, Ago2 as part of the RNA induced silencing complex (RISC), will be recruited and result in 3’ cleavage of the crRNA^19^. As miRNAs can be cell-specific, this synthetic construct would inactivate itself ubiquitously while only generating functional crRNA in a desired cell type where the cognate miRNA is present (Fig. 1a).

**Fig. 1:**
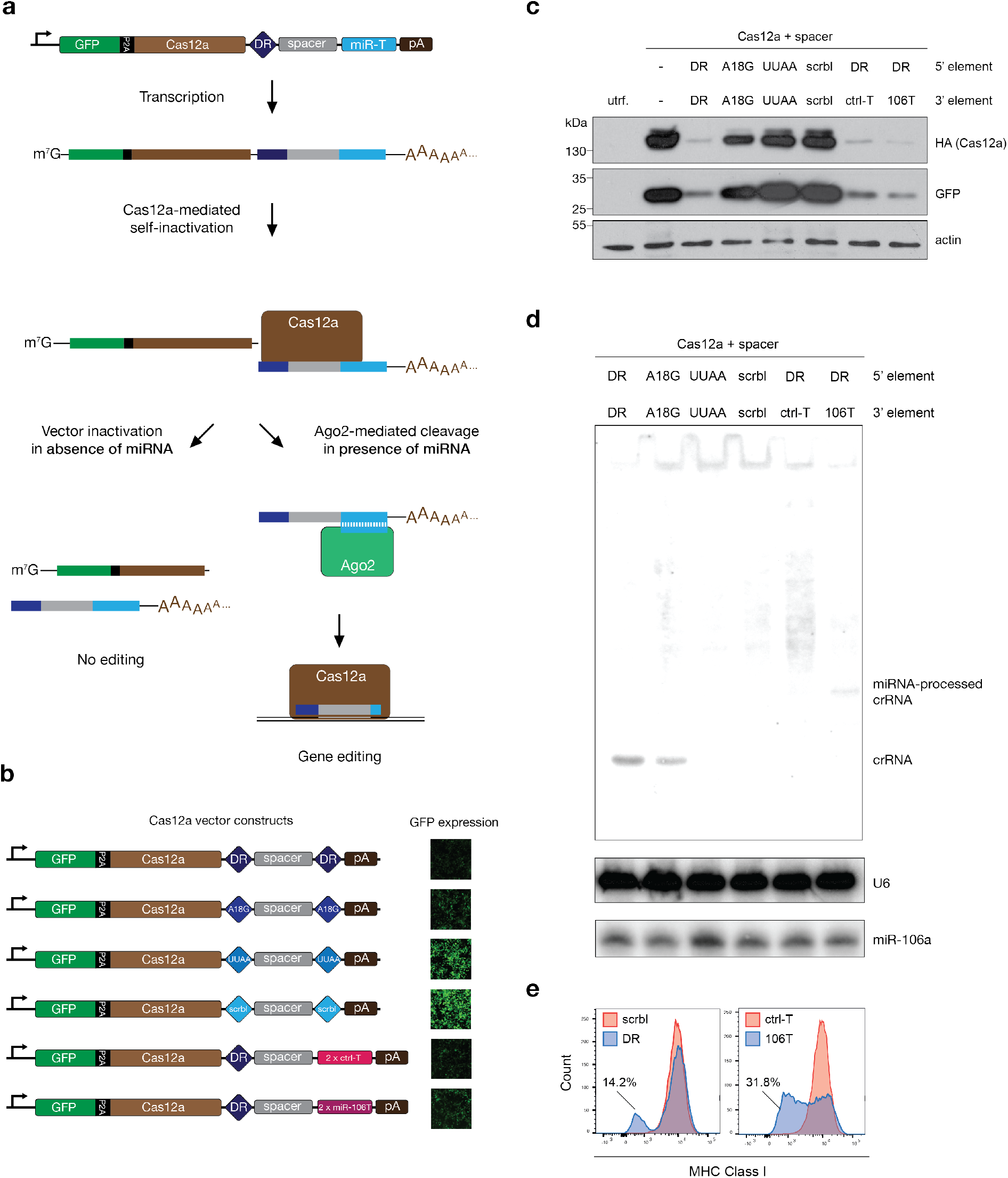
Self-inactivation and tissue specificity of Cas12a vector. **a,** A schematic of how self-inactivation and miRNA-based tissue specificity is incorporated into one single Cas12a vector. GFP and Cas12a are encoded in one open reading frame separated by a P2A site. The 3’-UTR of the same transcript consists of a direct repeat (DR), a spacer with complementarity to a genomic target and a sequence with perfect complementarity to a miRNA referred to as a miRNA-target site (miR-T) and lastly an SV40 polyadenylation signal (pA). **b**, Variations on the construct depicted in **a** with a mutation of nucleotide 18 in the direct repeat from an A to a G (A18G), inversion of nucleotides 16-19 from AAUU to UUAA (UUAA) or a scrambled sequence of the entire direct repeat (scrbl). The last two constructs have two target sites to either a control-miRNA (ctrl-miRT) or miR-106a (miR-106T). Fluorescence images are representative of GFP expression 48 hours after transfection with the constructs indicated. **c**, Western blot of cells transfected for 48 hours with the constructs depicted in **b**, -, no crRNA in 3’-UTR, utrf., untransfected cells. **d**, Small RNA Northern blot of total RNA from cells as in **c**, U6, U6 snRNA (loading control). **e**, MHC Class I cell surface expression measured ten days by flow cytometry ten days prost transfection. Data from cells transfected with the constructs overlaid as indicated.

To initially characterize self-inactivation, we experimented with variations on the DR design to ascertain the relationship between Cas12a engagement and loss of GFP expression (Fig. 1b). To this end, we utilized either canonical direct repeats, direct repeats that would be poorly or unable to be cleaved by Cas12a (A18G and UUAA, respectively), or one in which the direct repeats were disrupted altogether (scrambled; scrbl)^9,28^. To determine how these constructs would function, they were introduced into fibroblasts and monitored for GFP expression by both fluorescence microscopy and western blot (Fig. 1b-c). These data demonstrated that the GFP expression from the construct containing canonical direct repeats showed only low levels of fluorescence or expression by western blot which could also be correlated with HA-Cas12a expression. When the direct repeats were comprised of the A18G sites, fluorescence increased as compared to canonical sites. This enhanced expression could also be further corroborated by western blot analysis of both GFP and HA-Cas12a suggesting self-inactivation was diminished with the A18G sites. When the direct repeats were made to be uncleavable by Cas12a (UUAA), GFP expression was comparable to a construct lacking any direct repeats (scrbl). These data could be further validated by small RNA northern blot which indicated that wild type direct repeats, and to a lesser extent A18G, was capable of generating a visible crRNA, inversely correlating to the GFP and HA-Cas12a protein levels (Fig. 1c-d).

We next sought to make correct processing of the crRNA dependent on miRNA-mediated cleavage to enable cell-specificity. To this end, we replaced the 3’ direct repeat with two target sites for an endogenously expressed miRNA, miR-106a or an irrelevant control target sequence (ctrl-T) while keeping a wild-type direct repeat on the 5’ end to mediate self-inactivation (Fig. 1b). GFP and Cas12a protein expression from these two constructs were comparable to that containing two canonical direct repeats indicating that a single direct repeat is sufficient for self-inactivation (Fig. 1b-c). In contrast, the crRNA is no longer processed when the 3’ direct repeat is replaced with the control miRNA target sequence (ctrl-T) indicating a lack of cleavage (Fig. 1d). However, when ctrl-T is replaced with target sites corresponding to miR-106a, which is abundantly expressed in fibroblasts, we observe a distinct product corresponding to 16 Ago2-based cleavage (Fig. 1d). As Ago2 cleaves its target site between bases 10-11^16^, the resulting crRNA contains an extended 3’ terminus which should not impede Cas12a targeting or specificity^29^.

To ascertain whether the product of 5’ direct repeat and a 3’ miRNA cleavage site remains functional, we next expressed variants of our RNA construct that encoded a crRNA targeting beta 2 microglobulin (B2M). In comparing transcripts lacking direct repeats (scrbl), having both direct repeats, or containing a 5’ direct repeat with either a control 3’ target sequence (ctrl) or miR-106a 3’ sites we observe loss of MHC Class I, a proxy for B2M targeting, only in conditions in which the 3’ end of the spacer contained a wild type direct repeat or the miR-106a target sites (Fig. 1d). These data demonstrate a ~14% reduction of MHC1 with the canonical Cas12a targeting system which increases to greater than 30% targeting in the presence of miR-106a despite the extended crRNA (Fig. 1d-e). Together, these data suggest that miRNA biology can be exploited in conjunction with Cas12a-based processing to generate a single RNA capable of both self-inactivation and cell-specific targeting.

Given the capacity of a single transcript to both yield a functioning Cas12a editing platform and undergo self-inactivation, we next assessed whether this biologic circuit could be applied to a viral modality that would be most amenable to *in vivo* delivery. For safety reasons^30^, this work was done with an RNA virus incapable of spread and containing a scrambled crRNA with no complementarity to a genomic sequence. Utilizing only the RNA dependent RNA polymerase (RdRp) of Nodamura virus and the 5’ and 3’ uncoding material required for RdRp recognition as described elsewhere^31^, we generated a self-replicating RNA (herein referred to as a replicon) to express GFP, Cas12a, and a 3’ crRNA-containing UTR (Fig. 2a). As observed from our original plasmid design, launching of this replicon showed self-inactivation was also achieved using a viral-based delivery system (Fig. 2b). Comparable to what was observed using DNA, the RNA-based circuit equivalent showed self-targeting with canonical direct repeats (either one or two) with an intermediate phenotype observed for the A18G variant.

**Fig. 2:**
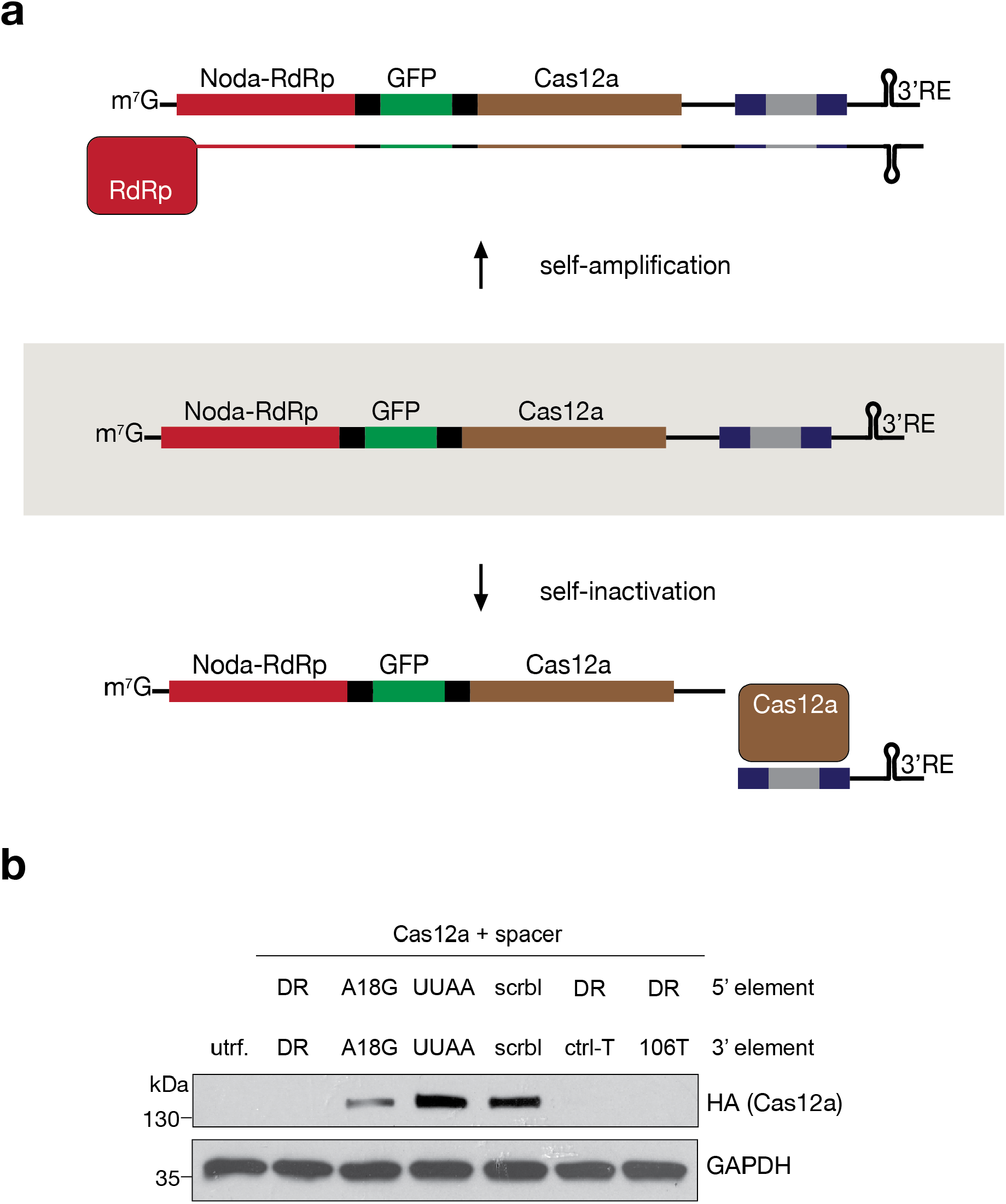
Delivery of Cas12a from a self-inactivating replicon. **a**, Schematic of a self-inactivating replicon construct. Nodamuravirus RNA dependent RNA polymerase (Noda-RdRp), GFP and Cas12a are encoded in one open reading frame separated by P2A sites. Downstream of the open reading frame, a spacer (grey) flanked by two direct repeats (dark blue) are inserted. **b**, Western blot of cells transfected with the replicon constructs indicated (as in Fig. 1c).

To determine whether the transcriptional response to our self-inactivating circuit would be amenable to *in vivo* use, we next performed bulk RNA sequencing to ascertain the transcriptional response to Cas12a expression and/or crRNA processing. To this end, we first compared the expression of Cas12a that was capable of self-inactivation and compared it to one incapable thereof. Surprisingly, this sequencing data set revealed that in contrast to sustained expression of Cas12a alone, the self-inactivating plasmid resulted in a significant number of differentially expressed genes (DEGs) (Fig. 3a and Supplementary Table 1). All upregulated genes with a log2fold change greater than 1 and an adjusted p-value less than 0.01 were annotated as belonging to the interferon response (Supplementary Table 1). These data would indicated that Cas12a processing of its own RNA results in a significant accumulation of aberrant RNA capable of inducing the host antiviral defenses. In contrast, this same comparison using the replicon-based platform yielded no DEGs (Fig. 3b). To determine if the lack of an interferon signature in response to the replicon-based platform was simply the result of having it generated in both conditions as a result of RdRp activity, we next compared the plasmid-based Cas12a system with processable crRNA to the equivalent replicon platform (Fig. 3c). This comparison yielded a larger number of DEGs but the interferon signature remained limited to plasmid-based delivery of Cas12a and crRNA demonstrating that the replicon self-inactivation is potent enough to prevent a cellular antiviral response (Supplementary Table 2). This was further corroborated by replicon read numbers which show that self-inactivation prevents any accumulation of either positive or negative sense transcripts that might otherwise serve as pathogen associated molecular patterns (Fig. 3d).

**Fig. 3:**
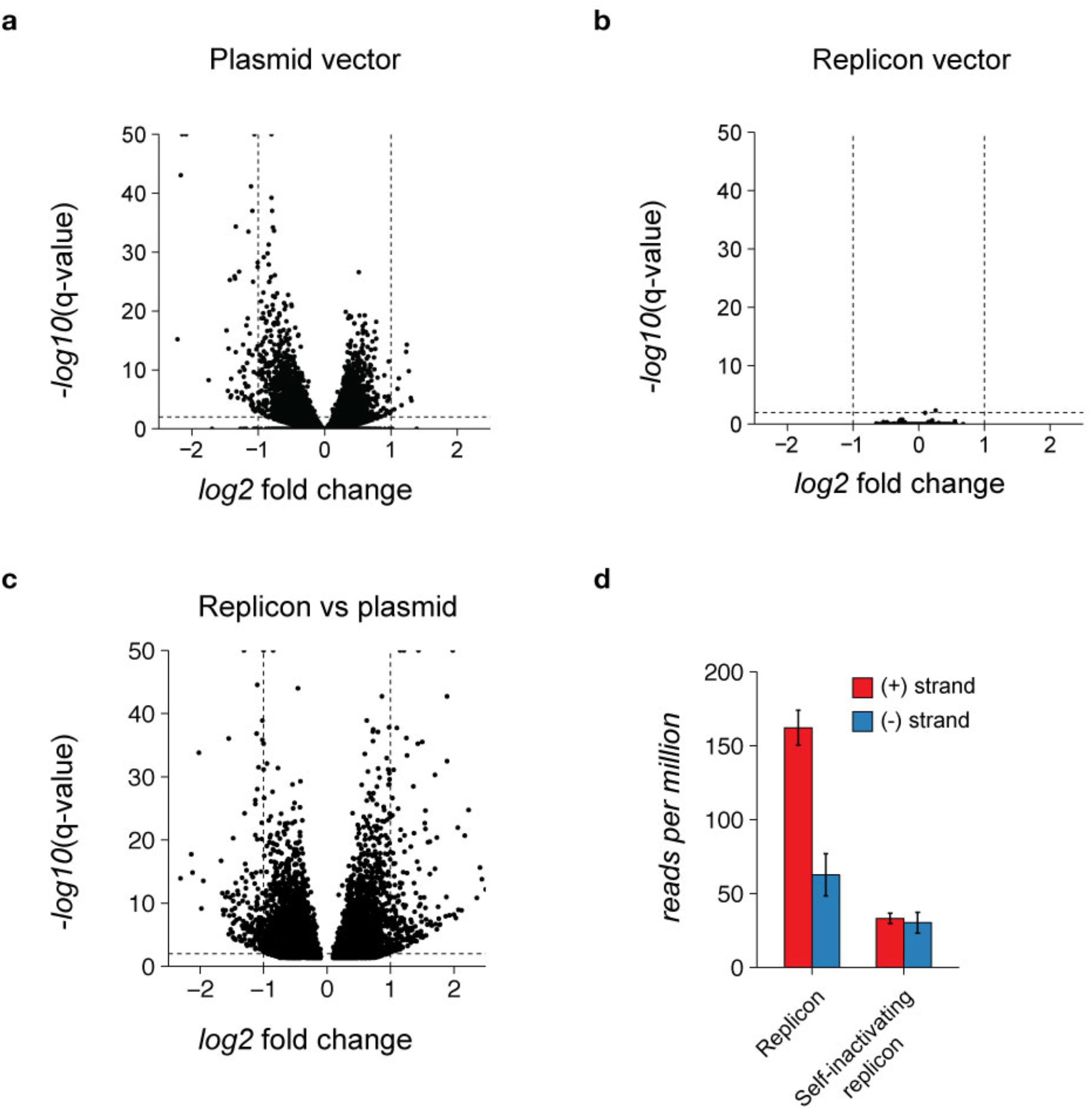
Transcriptional response to self-inactivating Cas12a vectors. **a**, Plot depicting differential gene expression of host genes in cells transfected with a plasmid-based Cas12a construct containing direct repeats in the 3’-UTR compared to cells transfected with a comparable construct without direct repeats. Each dot represents a gene plotted by its log2 fold change between the two conditions and -log10 of the adjusted p-value (q) determined based on triplicate samples. Horizontal line marks a q-value = 0.01 and Vertical lines mark a log2 fold change of −1 and 1. **b**, same as **a**, but comparing replicon-based Cas12a construct containing direct repeats against a comparable construct without direct repeats. **c**, same as **a**, but comparing plasmid-based Cas12a with direct repeats to replicon-based Cas12a with direct repeats. **d**, Stranded read numbers aligning to the replicon as number of reads per million of total reads. Error bars represent standard deviation from three replicates.

## DISCUSSION

Here we present data demonstrating that RNA-based platforms can be designed to support safe and effective genetic editing. Based on the dual RNase and DNase properties of Cas12a, we show that RNA constructs can be engineered to be self-targeting. This attribute ensures that Cas12a and crRNA expression is temporal, thereby minimizing off-target editing and the accumulation of aberrant and potentially inflammatory RNA. In addition, self-inactivation also allows one to adopt a RNA virus-based delivery system, as self-cleavage minimizes cytotoxicity and ensures no persistence or genomic integration.

Having shown that an RNA-based replicon can deliver a self-inactivated Cas12a and crRNA in the absence of a transcriptional response, we next sought to engineer this construct so that it would only function in a desired cell type. In general, nucleic acid-based therapeutics and gene therapy vectors rely on promoter elements that are uniquely specific to a desired cell type. While this strategy has achieved some noteworthy successes, use of DNA as a vector introduces other unwanted issues including the need for entry into the nucleus and the possibility of genomic integration. RNA-based vectors mitigate this risk by having no DNA phase and performing all of their function in the cytoplasm^32^. Given these attributes, we chose to adapt miRNA-based targeting as a means of instilling cell-specific activity. Here we show that the addition of a perfect complementary miRNA target site can replace the 3’ direct repeat needed to liberate a desired crRNA and thus render its biology cell specific. Together with the knowledge that every tissue or cell-type has a unique miRNA profile, these data suggest that one can engineer an RNA-based vector to efficiently enter the cytoplasm and then function only in those cells where editing is desired.

The completion of the human genome project ushered in hope that treating genetic diseases would soon become possible^33^. This dream was further fueled by the discovery and repurposing of the bacterial and archaeal CRISPR-Cas immune systems which provided unprecedented genome editing capabilities^1^. Indeed, recent efforts have suggested that this promise is closer to reality than ever before as demonstrated through the *ex vivo* editing of human T cells for therapeutic purposes^13^. While this latter accomplishment may provide countless genetic treatments, the full potential of CRISPR-based therapeutic still will require a vehicle for *in vivo* editing. Here we provide a platform to enable this next advancement. By exploiting the biology of RNA viruses, Cas12a, and miRNAs, here we demonstrate that one can design a single RNA that could be systemically delivered with high efficiency but only function for a set amount of time and only within a desired cell lineage. The basic principle underlying this biology could be used with the replicon based strategy outlined here or with existing RNA virus based genetic editors^32^. Taken together, with the rapid advancements of synthetic biology, new CRISPR-Cas systems, and our understanding of virus-host interactions, our progress towards *in vivo* editing may allow us to view genetic-based errors as something that can be seamlessly overwritten.

## METHODS

### Cells

All fibroblasts used in this paper were HEK293T cells which were maintained in Dulbecco’s Modified Eagle Medium (Gibco^®^) supplemented with 1x penicillin-streptavidin solution (Corning^®^) and 10% fetal bovine serum (Corning^®^).

### Plasmids

All constructs were synthesized by Twist Biosciences. The sequence will be deposited on NCBI and the plasmid will be made available on AddGene following publication.

### Western blot

Whole cell extract was prepared from live cells lysed in 1% NP-40 lysis buffer. 2μg (pEGFP plasmid-based experiments) and 70μg (nodamuravirus-based experiments) were analyzed by SDS-PAGE on a 4-15% acrylamide gradient gel (BIO-RAD^®^) and gels were subsequently blotted onto a 0.45μm nitrocellulose membrane (BIO-RAD^®^) and blocked in 5% milk in TBS for 1h at room temperature. Blots were probed with the following primary antibodies in 5% milk in TBS-T overnight at 4°C: anti-HA (clone HA-7, MilliporeSigma^®^), anti-GFP (ab290, Abcam^®^), anti-actin (clone Ab-5, Thermo Scientific^®^) and anti-GAPDH (G9545, MilliporeSigma^®^). After 4 x 5min washes in 1x TBS-T, blots were probed with HRP-linked secondary antibody for 1h at room temperature (anti-mouse, NA931V or anti-rabbit, NA934V, GE Healthcare^®^) and developed using the Immobilon Western HRP Substrate Kit (MilliporeSigma^®^).

### Small RNA Northern blot

Total RNA was extracted from live cells using TRIzol (Invitrogen). Northern blot was performed as described in with 20μg total RNA per sample^34^. Probes included the following: B2M-crRNA (5’-GCTGGATAGCCTCCAGGCCA-3’) miR-106a (5’-CTACCTGCACTGTAAGCACTTTT-3’) and U6 (5’-GCCATGCTAATCTTCTCTGTATC-3’). Probes were labeled with ATP-P32 using T4 polynucleotide kinase (NEB^®^) and blot was exposed to a phosphor screen (GE^®^) and developed on a Typhoon TRIO Storage Phosphorimager (GE^®^).

### Flow cytometry

Roughly 7.5×10^5^ cells/well were plated on 6-well plates. After attaching overnight, cells were transfected using lipofectamine 2000 (Invitrogen^®^) and were passaged 1:5 when they reached ~80% confluency for up to ten days. For flow cytometry analysis, cells were trypsinized and washed 2 x stain buffer (FBS) (BD Biosciences^®^) and stained using the BD Cytofix/Cytoperm Fixation/Permeabilization Kit (BD Biosciences^®^). The following antibodies and dyes were used: anti-human HLA-A,B,C Pacific Blue (clone W6/32, BioLegend^®^), anti-HA Alexa Fluor 647 (clone HA.11, BioLegend) and LIVE/DEAD stain Aqua (ThermoFisher^®^). Fixed cells were analyzed on a 2019 Attune NxT Flow Cytometer. Data processing was done with FlowJo v. 10.6.2.

### RNA sequencing

Total RNA was extracted using TRIzol (Invitrogen^®^). 1μg of total RNA per sample was used as starting material for building Illumina libraries for deep sequencing. Libraries were built following Illumina protocols using the TruSeq Stranded mRNA library prep kit (Illumina^®^). Libraries were enriched for polyadenylated transcripts and sequenced on an Illumina NextSeq 500. Alignment of reads was done using STAR alignment to the human reference genome (hg38) and differential gene expression and statistics were determined by the DEseq2 protocol. Reads mapping to the Nodamuravirus replicon were aligned using Bowtie2.

## SUPPLEMENTARY MATERIAL

**Table 1. Differentially expressed genes in response to self-inactivating vs. non-processable plasmid-based Cas12a vector**. The control group are fibroblasts expressing GFP, Cas12a, and a crRNA lacking direct repeats performed in triplicate. The experimental group are fibroblasts expressing GFP, Cas12a, and a crRNA flanked with direct repeats, also performed in triplicate. Total reads per sample are given. Shown are gene names, mean count per gene, log2fold change in comparing triplicate samples in deSeq2, Standard error and adjusted p-value (q value). Raw data to be deposited on NCBI GEO.

**Table 2. Differentially expressed genes in response to self-inactivating plasmid-based Cas12a versus the corresponding replicon-based construct. The control group are fibroblasts expressing GFP, Cas12a, and a crRNA flanked with direct repeats performed in triplicate. The experimental group are fibroblasts expressing GFP, Cas12a, and a crRNA flanked with direct repeats as delivered by RNA replicon, also performed in triplicate. Total reads per sample are given. Shown are gene names, mean count per gene, log2fold change in comparing triplicate samples in deSeq2, Standard error and adjusted p-value (q value). Raw data to be deposited on NCBI GEO.**

